# Mycelium of mushroom-producing fungi as high-quality protein source

**DOI:** 10.1101/2025.07.02.662711

**Authors:** Jasper Zwinkels, Philip van de Koolwijk, Nikkie van der Wielen, Oscar van Mastrigt, Eddy J. Smid

**Affiliations:** Food Microbiology, Wageningen University, Wageningen, the Netherlands; Animal nutrition, Wageningen University, Wageningen, the Netherlands

## Abstract

To achieve a more sustainable food system, reducing reliance on animal-based proteins is essential. Mycoprotein from mushroom-producing fungi (Basidiomycota) presents a promising yet underexplored alternative. Basidiomycetous mycelium combines the high protein quality of fungi with the sensory qualities of mushrooms. This study compares the mycelium of commercially cultivated and edible basidiomycetes to their fruiting bodies and to mycelium of *Rhizopus microsporus* var*. oligosporus* (tempeh fungus). Nutritional quality was assessed via protein content and PDCAAS, while sensory potential was analysed through equivalent umami concentration (EUC) and pyrazine levels. High-performing species surpassed the EUC of *R. microsporus* and fruiting bodies, reaching 301.0 g MSG-eq/100 g DW. Basidiomycetous mycelia exhibited superior protein quality (PDCAAS up to 1.20) and utilizable protein content (33.2 g/100 g DW), qualifying as an “Excellent protein source.” Given the phylogenetic diversity of basidiomycetes, this study highlights the untapped potential of its mycelium as a high-quality protein source.

## 1 Introduction

Our current food production system exerts considerable environmental pressure, largely due to livestock herding for meat production, which accounts for 57% of anthropogenic greenhouse gas (GHG) emissions (Crippa et al., 2021; Xu et al., 2021). This has driven increasing interest in sustainable alternatives to animal protein. One promising option is mycoprotein, a mycelium-based protein from filamentous fungi. Particularly, mycelium from mushroom-producing fungi (basidiomycetes) could be a promising source, combining benefits of mushrooms and mycelia. A recent study showed that replacing just 20% of ruminant meat with mycoprotein could halve related deforestation and GHG emissions, highlighting the large potential impact of shifting toward fungal proteins (Humpenöder et al., 2022).

Despite their environmental benefits, fungal-based foods, including mycoprotein, have not yet been widely accepted. This slow adoption is influenced by multiple factors, such as flavour and health, familiarity, attitudes, food neophobia and social norms (Onwezen et al., 2021), with taste and health being particularly influential (Jahn et al., 2021; Schmidt & Mouritsen, 2022). One aspect of the widespread appeal of meat is its umami taste and meaty-roasted aromas, with umamibeing identified as a key driver of consumer acceptance (Miller et al., 2014; Schmidt & Mouritsen, 2022). These sensory characteristics often remain absent in meat alternatives and enhancing these could significantly improve their likeability and acceptance.

Umami taste is primarily mediated by the synergistic action of free amino acids and 5’-nucleotides, sensed via specific taste receptors (Yamaguchi et al., 1971). Their combined effect is often expressed as equivalent umami concentration (EUC), which correlates strongly with perceived umami intensity (Zhu et al., 2022). Particularly mushrooms are meat alternatives that are known and appreciated for their high umami content(Liu et al., 2012; Mau, 2005). Little is known about the umami properties of mycelial products, such as mycoprotein or fungal-fermented foods like tempeh, but there are indications that the mycelium of mushroom-producing fungi could contain similarly high levels of umami compounds (Liu et al., 2012; Mau, 2005).

In addition to taste, aroma plays a critical role in the sensory appeal of meat alternatives. Pyrazines, a class of nitrogen-containing aromatic compounds, are central to the characteristic scent of cooked meat, contributing roasted, nutty, and beef-like notes (Christlbauer & Schieberle, 2009; Moon et al., 2006). These compounds are also abundant in mushrooms (Zhang et al., 2018; Zhuang et al., 2020), making fungal proteins a promising alternative not only for nutrition but also for their aromatic properties.

In addition to sensory aspects, consumers and experts have raised concerns about adequate protein intake with reduced consumption of meat, due to the generally lower protein content and quality in alternative protein sources (Moughan, 2021; Onwezen et al., 2021; Tso & Forde, 2021). For example, mushrooms exhibit a protein digestibility-corrected amino acid score (PDCAAS) of 0.2–0.7, compared to 0.92–1.17 in meats (Ayimbila & Keawsompong, 2023; Dabbour & Takruri, 2002; Fanelli et al., 2022). Many other meat alternatives, such as plant-based proteins, face similar limitations in terms of protein quality (Erbersdobler et al., 2017; Ertl et al., 2016; Fanelli et al., 2022; Negrão et al., 2005; Zwinkels et al., 2023). These limitations raise concerns about their ability to support human protein requirements.

In contrast, mycelium-based proteins like mycoprotein typically have higher protein quality. Quorn, a commercially available mycoprotein made from *Fusarium venenatum*, has a PDCAAS of 0.996 (Edwards & Cummings, 2010), comparable to that of other conventional food fungi (CFF) such as *Rhizopus microsporus* var. *oligosporus*, the most consumed edible fungus globally (Grand view research, 2024; R. Wang et al., 2023).

These considerations suggest that the mycelium of mushroom-producing fungi may offer a compelling new protein source, combining the nutritional benefits of CFF with the desirable sensory characteristics of mushrooms. To harness the potential of the large and diverse phylum of basidiomycetes, it is essential to evaluate how the nutritional quality, taste, and aroma of basidiomycetous mycelia compare to those of their fruiting bodies and the mycelium of CFF.

Therefore, this study aims to evaluate the potential of basidiomycetous mycelia as a protein source for human consumption by comparing mycelia of phylogenetically diverse edible basidiomycetes to their fruiting bodies and to the mycelium of a CFF (*R. microsporus*). We assess taste and aroma, through umami-active compounds and alkylpyrazine content, as well as nutritional value, measured via protein content and protein quality.

## 2 Results

### 2.1 Taste & aroma of mycelium and fruiting bodies

#### 2.1.1 Equivalent umami concentration highly varied between species

The umami taste of basidiomycetous mycelium (myc) of eight species was compared to five corresponding species of fruiting bodies (FBs) and *R. microsporus* myc through the equivalent umami concentration (EUC) (**Fig. 1**). The EUC highly varied in basidiomycetous myc between species, ranging from 11.9-301.0 g MSG-eq/100 g DW, but in general was significantly higher than in FBs (t-test, p=0.01) (26.8-94.9 g MSG-eq/100 g DW) and *Rhizopus* myc (23.9 g MSG-eq/100 g DW). The five highest EUC values were observed in basidiomycetous myc *(S. commune, P. pulmonarius, F. velutipes, S. rugoso-annulata, V. volvacea*), reaching level 2 EUC (100-1000 g MSG-EQ/100 g DW), compared to level 3 (10-100 g MSG-EQ/100 g DW) in other samples, as defined by Mau (2005).

**Fig. 1.**
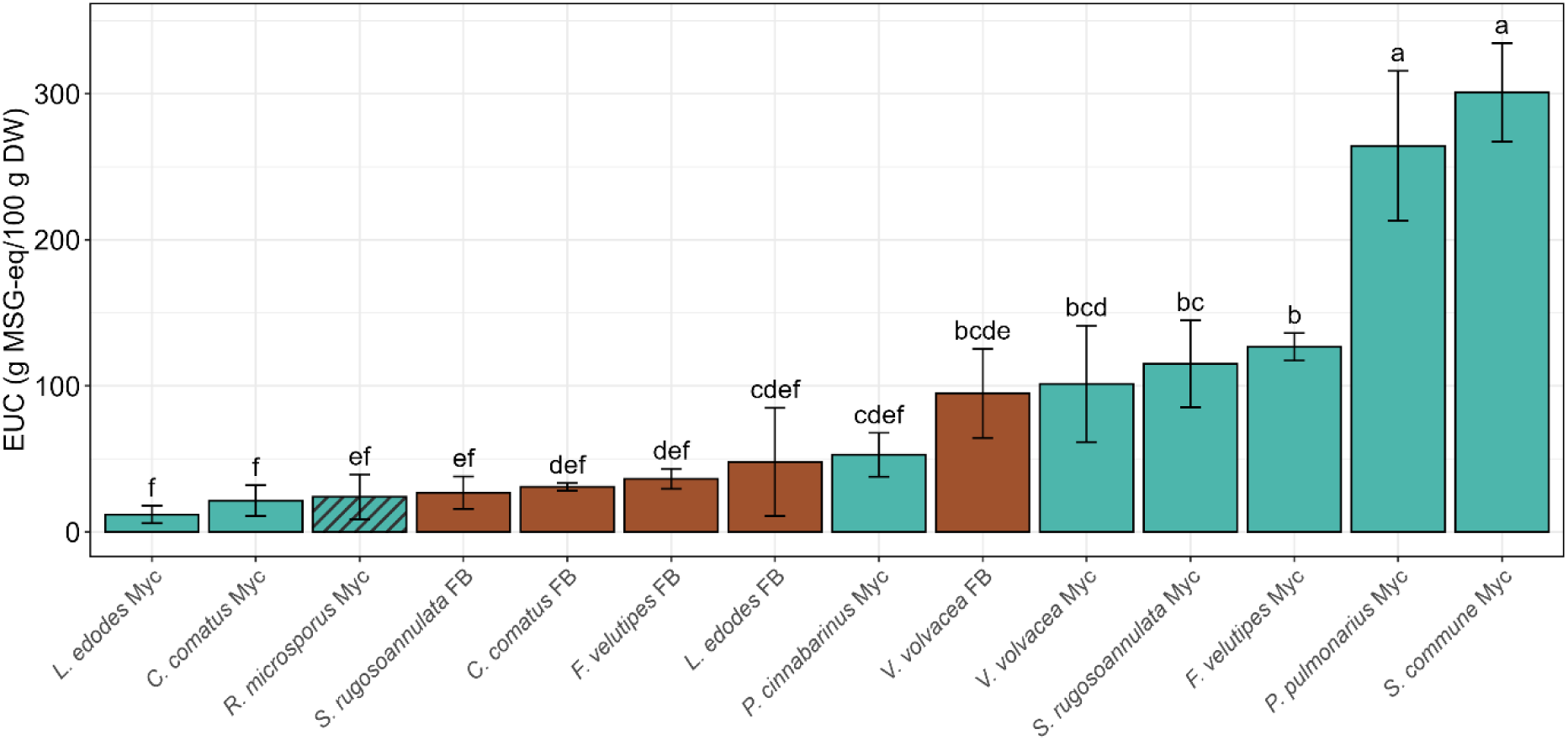
Equivalent umami concentration (EUC) of mycelium and fruiting bodies. EUC of basidiomycetous mycelium (cyan), fruiting bodies (brown) and *Rhizopus* mycelium (dashed cyan). Bars are means of biological triplicates ± standard deviation. Different letters express statistical significance (TukeyHSD, p<0.05)

Among the four nucleotides measured, 5’-guanosine monophosphate (GMP) dominated the contribution of nucleotides to the EUC, while among the umami FAA glutamate was the main contributor (Supplementary fig. 1). The sweet and bitter-tasting FAA likely contributed far less to the taste of fungal materials than umami FAA, expressed in their lower dose over threshold (Supplementary fig. 2).

#### 2.1.2 Mushroom alcohols are also present in mycelium

Mushroom alcohols are important aroma compounds in mushrooms and meat, providing an earthy, fungal aroma and improve saltiness perception (Xiao et al., 2024). Mushroom alcohols (1-octen-3-ol, 3-octanol, 2-octen-1-ol, and 3-octen-2-one) were formed in raw mycelium, fruiting bodies, and beef (**Fig. 2**a). During heating, mushroom alcohols remained unchanged (t-test, p > 0.05, Supplementary fig. 3). The mushroom alcohol with the highest abundance was 1-octen-3-ol, which was found in six basidiomycetous mycelia, two fruiting bodies, and beef (**Fig. 2**b&c).

**Fig. 2.**
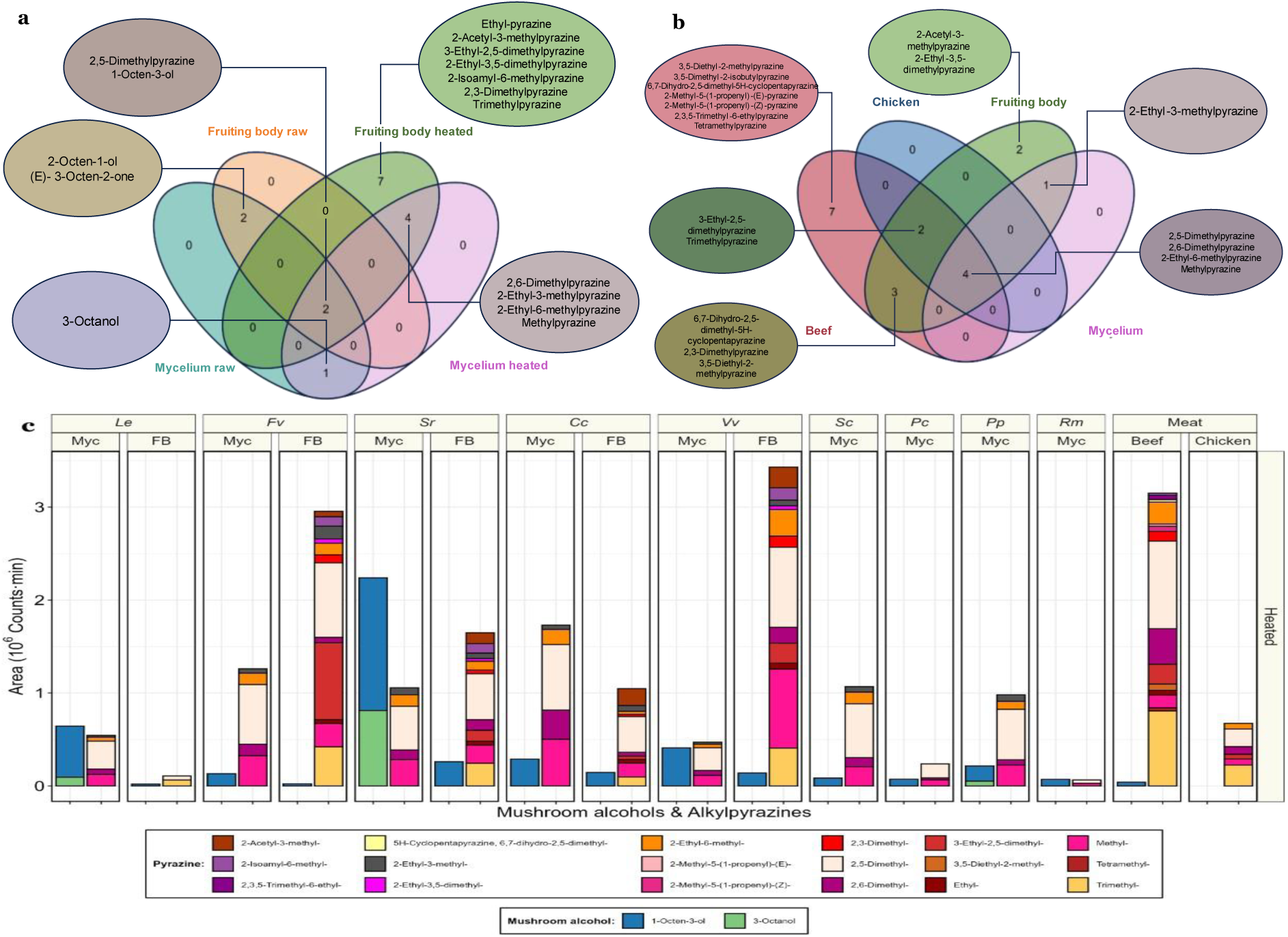
Mushroom and roasted meat-like volatile compounds are present in mycelium and fruiting bodies. a) Venn diagram of raw mycelium (turquoise), raw fruiting body (orange), heated fruiting body (green) and heated mycelium (pink). Values indicate the number of compounds in a certain group, while the names allude to mushroom alcohols and pyrazines of interest. b) Venn diagram of heated beef (red), heated chicken (blue), heated fruiting body (green) and heated mycelium (pink). Numbers represent the number of pyrazines present in a group. Area over 20.000 counts*min was determined as present. c) Mushroom alcohol (left) and alkylpyrazine (right) area (106 counts*min) in heated mycelium and fruiting bodies and chicken and beef. Species are *L. edodes* (*Le*), *F. velutipes* (*Fv*), *S. rugosa-annulata* (*Sr*), *C. comatus* (*Cc*), *V. volvacea* (*Vv*), *S. commune* (*Sc*), *P. cinnabarinus* (*Pc*), *P. pulmonarius* (*Pp*), and *R. microsporus* (*Rm*). Myc = Mycelium, FB = Fruiting body. Colours represent compounds and are explained in the figure legend.

#### 2.1.3 Pyrazines are formed during heating in mycelium and fruiting bodies

Pyrazines are aromatic nitrogenous compounds commonly found in mushrooms and known for their contribution to the aroma profile of meat, typically imparting roasted, nutty, or meaty notes (J. Li et al., 2024; Z. Li et al., 2021). In raw material, only 2,5-dimethylpyrazine was detected. However, upon heating, the number of unique alkylpyrazines increased, with mycelium containing five, fruiting bodies containing twelve, and beef containing sixteen different pyrazines. Furthermore, the total area significantly increased by 27-85-fold (when also detected in raw material), from a maximum of 6.1*10^4^ in raw fungi to 6.5*10^5^-3.4*10^6^ counts*min in heated fungi (**Fig. 2**c, Supplementary fig. 3). The lowest total area in heated material was observed in *R. microsporus* myc (6.6*10^4^ counts*min), while basidiomycetous myc contained significantly more alkylpyrazines (t-test, p<0.01), reaching similar levels as in chicken (t-test, p>0.05). However, on average basidiomycetous myc contained significantly less alkylpyrazines than their fruiting bodies (t-test, p<0.05). *V. volvacea* FB (3.4*10^6^ counts*min) and *F. velutipes* FB (3.0*10^6^ counts*min) contained alkylpyrazine levels in the range of beef (3.2*10^6^ counts*min). Despite the higher diversity of alkylpyrazines in beef, the unique compounds only amounted to 1% of its total area, thereby likely contributing little to the overall aroma.

Furthermore, pyrazines have varying, but related odour descriptions and thresholds, ranging from roasted beef and nutty, to burnt and popcorn (Supplementary table 1). The most abundant compound in heated meat and fungal material was 2,5-dimethylpyrazine, contributing to 28-30% of the total area in meat, and 25-55% in fungal material. The odour of 2,5-dimethylpyrazine is described as roasted beef, cocoa, and nutty. The alkylpyrazines detected had vastly different odour thresholds, ranging from 0.002 to 100,000 µg/L (Supplementary table 1). Therefore, consideration of the odour activity value (OAV), calculated by the dividing the area by the odour threshold (OT), might be a better metric to estimate the aroma perception of the compound. The OAV showed an inflated gap between fruiting bodies, mycelium, and meat, although differences between fruiting bodies and beef remained insignificant (Supplementary fig. 4, LMM, p>0.05). Additionally, the AOV of basidiomycetous mycwas significantly lower than chicken, but remained significantly higher than *Rhizopus* mycelium (LMM, p<0.01).

### 2.2 Protein content is higher in mycelium than fruiting bodies

Nitrogen content of mycelium and fruiting bodies was determined by DUMAS combustion method and converted to crude protein (CP) content using a fungi-specific conversion factor (Mariotti et al., 2008). Crude protein contents ranged from 17.7 g/100 g DW in *C. comatus* fruiting body to 35.9 g/100 g DW in *R. microsporus* myc (Fig. **3**). The CP content of *S. rugosa-annulata* and *S. commune* mycelia was comparable to that of *R. microsporus*, while other basidiomycetous mycelia had significantly lower contents. Despite this, all mycelia contained higher CP contents than their fruiting bodies, up to 1.7-fold, and significantly higher in three species.

**Fig. 3.**
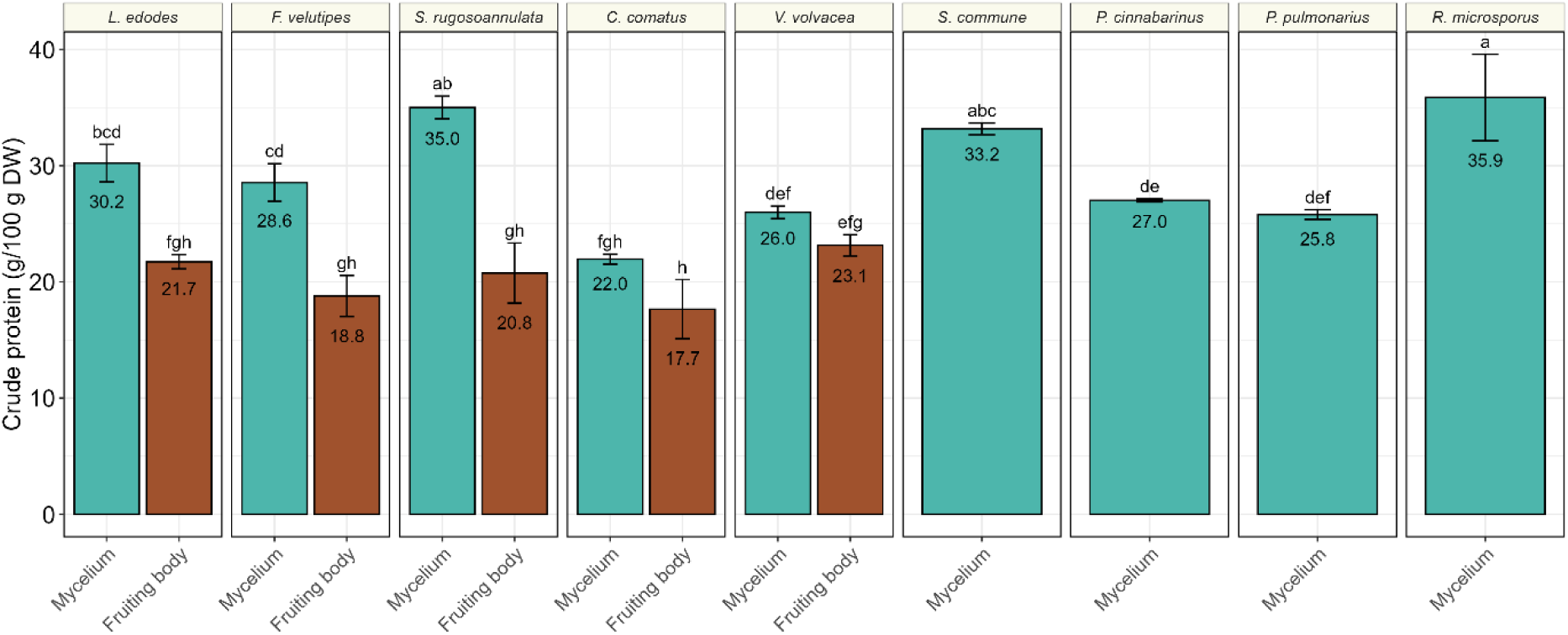
Crude protein content of mycelium and fruiting bodies. Red bars represent mycelium, blue bars the fruiting bodies. Bars are means of biological triplicates ± standard deviation. Numbers indicate the mean crude protein content. Different letters indicate a significant difference (TukeyHSD, p<0.05).

### 2.3 Protein quality is highest in basidiomycetous mycelium

Protein quality in mycelia of basidiomycetes was assessed by the protein digestibility corrected amino acid score (PDCAAS) and compared to their fruiting bodies and mycelium of *R. microsporus*. PDCAAS was calculated based on the amino acid score of the lowest amino acid, which was based on the dietary requirements of a pre-school child (0.5-3 yr.) and corrected for the *in vitro* protein digestibility (IVPD) of the sample (FAO, 2013). The IVPD of basidiomycetous mycelia was high (90.2-100.0%) (Supplementary fig. 6) and significantly higher than the corresponding fruiting bodies (ANOVA, p<0.0001), up to 12.6% in *V. volvacea*. Similarly, the IVPD of basidiomycetous mycelia was significantly higher than of *R. microsporus* (77.9%; ANOVA, p<0.0001), with *P. cinnabarinus* reaching the highest value (1.20). Four of eight species of basidiomycetous mycelia surpassed a PDCAAS of 1.0. Additionally, the amino acid composition in basidiomycetous mycelia was of higher nutritional quality, than in fruiting bodies and *R. microsporus* myc. Firstly, this is indicated by the higher indispensable amino acid index the mycelium (Supplementary table 2). Secondly, four out of five fruiting bodies and *Rhizopus* myc did not meet dietary requirements, and were limited in either leucine, lysine, sulphur amino acid (SAA), or tryptophan (trp) (Supplementary table 3). In contrast, six out of eight basidiomycetous mycelia had a complete amino acid profile without any limiting amino acids.

Utilizable protein represents the amount of protein that can effectively be absorbed, and used by the body for growth, maintenance, and physiological functions. It was calculated by multiplying the truncated PDCAAS by the crude protein content. **Fig. 5** illustrates the relationship between the utilizable protein content and the equivalent umami concentration (EUC), integrating taste potential, protein content and protein quality. The utilizable protein content of all mycelia (20.5 – 32.6 g/100 g DW) was significantly higher than in fruiting bodies (11.2 – 19.6 g/100 g DW) (LMM, p< 0.0001; Supplementary fig. 7). Among all samples, mycelia of *S. rugosa-annulata* and *S. commune* had the highest utilizable protein contents (32.6 and 33.2 g/100 g DW, respectively), both significantly exceeding that in *R. microsporus* myc (27.3 g/100 g DW). Conversely, *C. comatus*, and V. *volvacea* had the lowest utilizable protein contents in basidiomycetous mycelia (20.5 and 21.3 g/100 g DW, respectively), which were significantly lower than in *R. microsporus*. When combining utilizable protein content and EUC, mycelia of the basidiomycetes *V. volvacea*, *P. pulmonarius*, *F. velutipes*, *S. rugosa-annulata*, and *S. commune* outperformed fruiting bodies, of which the latter three outperformed *R. microsporus* myc, and *S. commune* myc ranking the highest in both parameters.

**Fig. 4.**
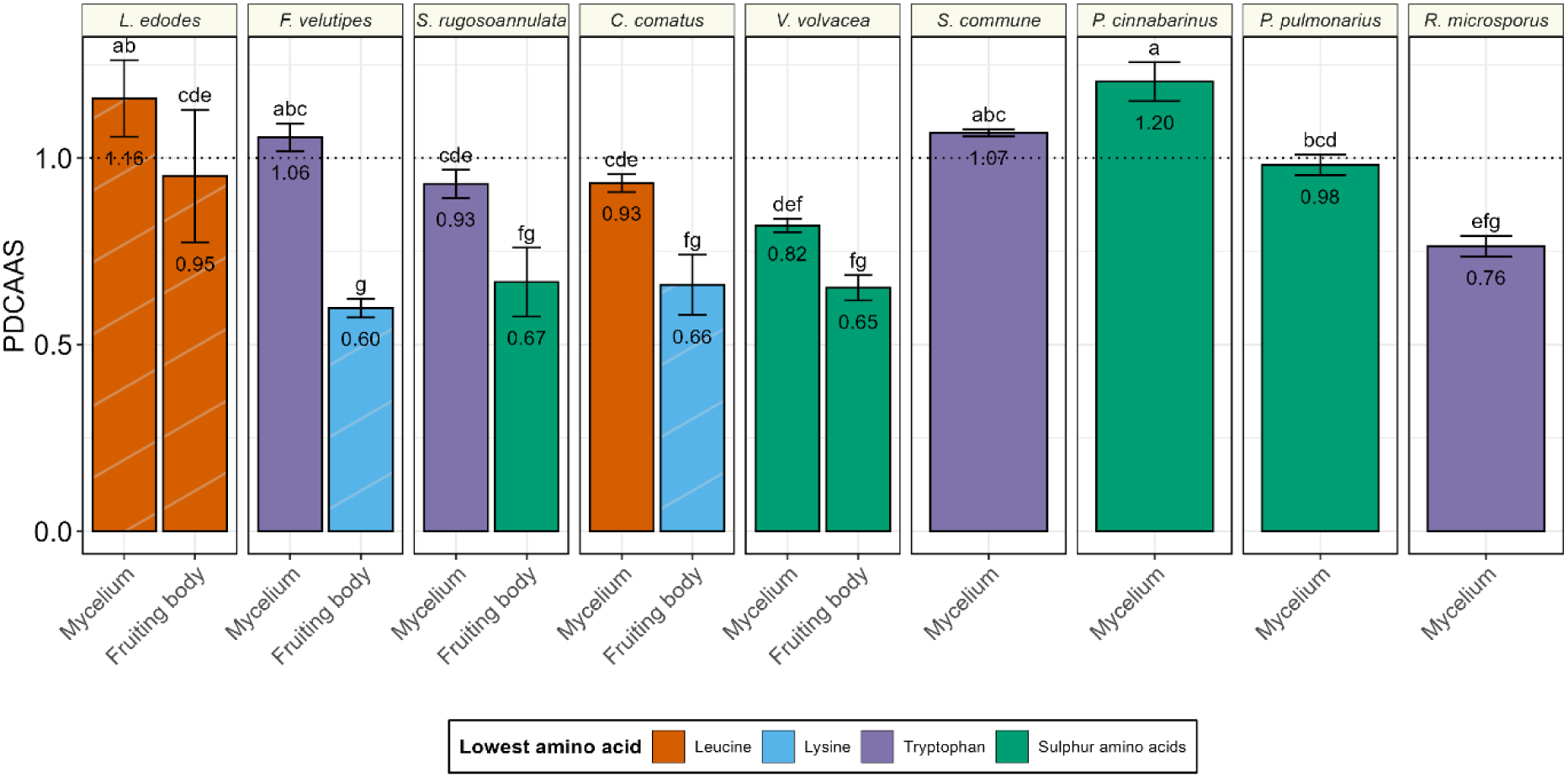
Protein digestibility corrected amino acid score (PDCAAS) of mycelium and fruiting body of basidiomycetes and *Rhizopus*. Bars are means of biological triplicates ± standard deviation. Colours indicate the lowest amino acids leucine (red), lysine (green), sulphur amino acids (SAA, blue), and tryptophan (purple). Lowest amino acid is the limiting amino acid when PDCAAS < 1.0 (dotted line). SAA (cysteine and methionine) and tryptophan were not measured in *L. edodes* mycelium and fruiting body, *F. velutipes* fruiting body, and *C. comatus* fruiting body (stripes). Different letters indicate a significant difference (TukeyHSD, p<0.05).

**Fig. 5.**
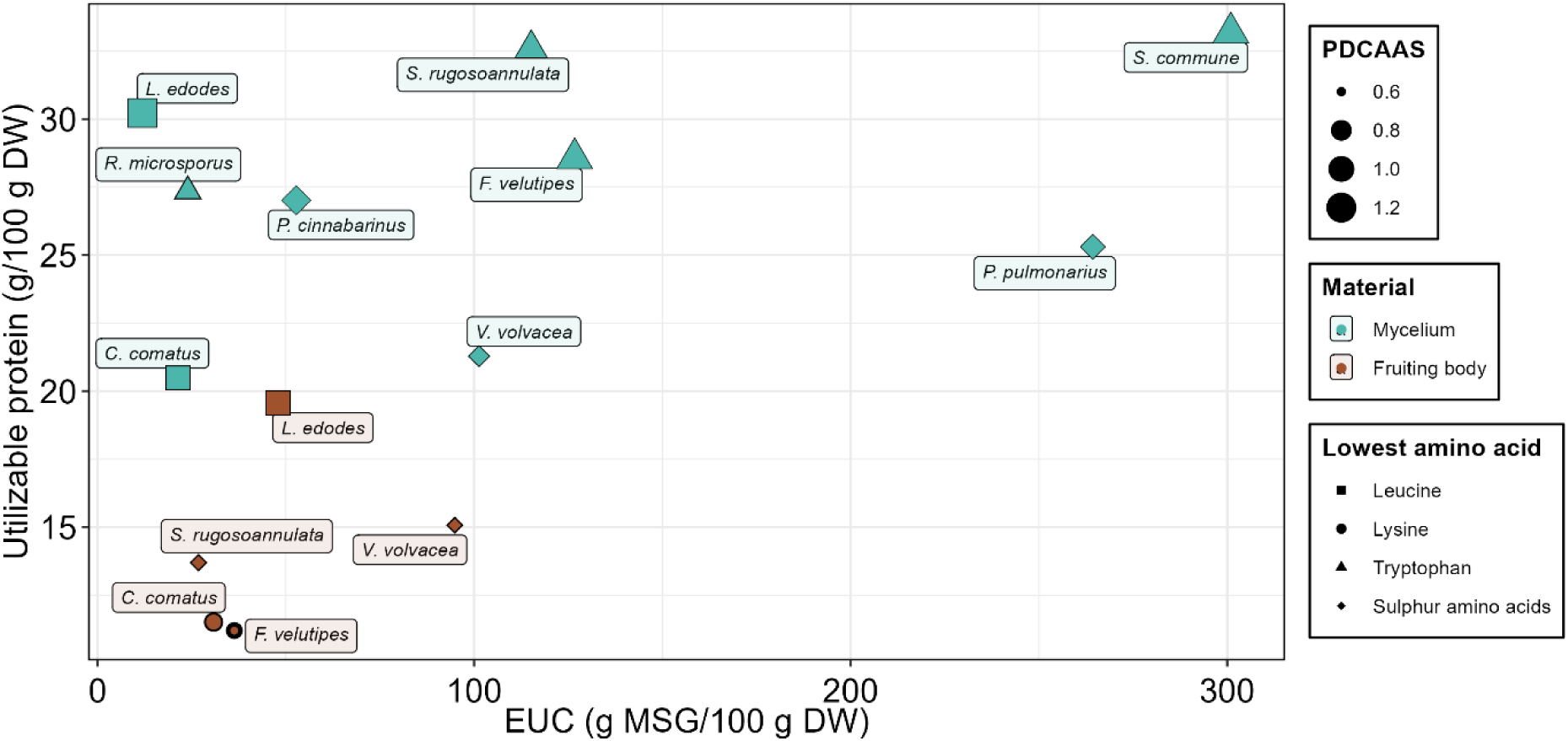
Mycelium of Basidiomycetes outperforms *R. microsporus* mycelium and mushroom fruiting bodies in terms of protein quality and umami taste. Scatter plot of mycelia (red) and fruiting bodies (blue) based on utilizable protein content (g/100 g DW) and equivalent umami concentration (g MSG-EQ/100 g DW). Point shapes represent the lowest amino acid, while point size corresponds to the PDCAAS.

## 3 Discussion

This study explored the taste, aroma, and nutritional qualities of mycelium of eight species from the phylum of Basidiomycota and compared these to the fruiting bodies of five corresponding species and the mycelium of *Rhizopus microsporus* var. *oligosporus*, by analysing the content of umami-active compounds, alkylpyrazines, and the protein content and quality. With the assessment of these parameters, we aimed to demonstrate the potential of the mycelium of basidiomycetes as a novel protein source for human consumption.

Umami taste is a critical factor in low-sodium food likeability and novel food acceptance (Miyaki et al., 2016; Schmidt & Mouritsen, 2022) and mushrooms generally have a high umami taste and EUC, which contributes to their appreciation as protein source (Pashaei et al., 2024). Moreover, umami taste enhances saltiness perception, which can increase food acceptability at reduced salt levels (Ma et al., 2024). The addition of umami compounds enables a 30% reduction in salt while maintaining the same acceptability (Rosa et al., 2021). Substitution of meat with mushrooms in reduced salt formulations even maintains acceptability at lower salt perception levels (Miller et al., 2014). This is an important benefit, considering that salt intake is a major risk factor for human health in a Western diet and plant-based meat alternatives generally contain a lot of salt (Nájera Espinosa et al., 2024; Q. Yang et al., 2024). In addition to taste, salt is often added as a preservative. However, even at low salt concentrations, fungal foods such as tempeh and Quorn have substantial shelf lives up to several weeks.

This study demonstrates that high EUC, which correlates highly with umami taste, is also present in the mycelium of basidiomycetes. Three out of five species contained even higher levels of EUC in the mycelium than in their fruiting bodies. And the five highest ranking samples were mycelia. These five mycelia were the only ones to reach umami level 2 (Yamaguchi et al., 1971), thereby coming close to the EUC observed in sous-vide cooked beef (Hwang et al., 2020). Even though a high umami taste is generally observed in fruiting bodies, a higher EUC in mycelia has been observed before in several Tuber species (Liu et al., 2012).

However, in some species fruiting bodies contained higher EUC levels than the mycelium, such as in *L. edodes* and *C. comatus*, which contained 4 and 1.4-fold higher EUC than mycelium, in line with even greater differences found in literature (sixfold and 25-fold, respectively) (Tsai et al., 2007a; W. K. Wang et al., 2016). Notably, the EUC in fruiting bodies can vary, depending on growth and storage factors, which includes some inherent variation in these comparisons (Ren et al., 2022; Shi et al., 2025; Tsai et al., 2007; J. Wang et al., 2023; Xia et al., 2021; J. H. Yang et al., 2001; Zou et al., 2023).

Compared to other mycelium, mycelium of basidiomycetes contained up to 12.5 times higher EUC than *R. microsporus.* This indicate that the mycelium of specifically basidiomycetes is well equipped to accumulate umami FAA and 5’-nucleotides. The reason for the accumulation of these compounds in basidiomycetes could lie in the developmental cycle of this phylum. Namely, Son et al. (2019) suggest that the accumulation of FAA and nucleotides starts the development of fruiting bodies from mycelia. This implies that high levels of these compounds are typically associated with well-developed mycelium and newly formed mushrooms, which has implications for the harvest time of mycelia to optimize umami taste.

Lastly, the dose over threshold (DOT) of umami FAA was generally 2 to 3 orders of magnitude higher than of neutral, sweet, and bitter FAA (Supplementary fig. 2). This indicates that overall FAA likely contributed more to the umami taste of the mycelium and fruiting bodies than bitter or sweet taste.

Mushrooms have a distinct aroma profile, partly arising from compounds called mushroom alcohols. Mushroom alcohols are also found in meat and enhance flavour by increasing saltiness (Xiao et al., 2024). Therefore, their presence in mycelia could enhance their palatability. The main mushroom alcohol, 1-octen-3-ol (Venkateshwarlu et al., 1999), was formed in raw mycelia and its abundance remained stable during heating, ensuring its contribution to the aroma profile in cooked mycelium. Interestingly, 1-octen-3-ol was significantly more abundant in basidiomycetous mycelium than in *R. microsporus* myc and most fruiting bodies, which may reflect a more favourable aroma profile of basidiomycetous mycelium relative to CFF.

Similarly, alkylpyrazines are important compounds in the aroma profile of roasted meat, mushrooms, and fermented foods like natto, and soy sauce (Moon et al., 2006; Rocchi et al., 2024; Zhang et al., 2018). They are generally formed during Maillard reaction in low water environments (Adams & Kimpe, 2009; Amrani-Hemaimi et al., 1995). 2,5-dimethylpyrazine was likely produced during fermentation, since it was the only alkylpyrazine present in raw fungal material. Other alkylpyrazines were likely a result of the Maillard reaction occurring during heating, similarly to meat. However, raw mycelium does contain the precursors for alkylpyrazine production, in the form of free amino acids and in the raw materials. In this study, beef contained the highest variety in alkylpyrazines, followed by mushrooms, chicken, and mycelium. However, beef is generally more diverse in alkylpyrazines than other meats (Sohail et al., 2022). Heated basidiomycetous mycelium contained a lower area and AOV than fruiting bodies and beef, but similar to chicken, and higher than *R. microsporus* myc. Therefore, basidiomycetous mycelia might have a stronger roasted, meat-like aroma than CFF, improving their palatability, although not to the extent found in mushrooms.

In addition to the flavour, protein content analysis revealed that basidiomycete myc is rich in protein, with crude protein levels significantly exceeding those of their fruiting bodies. Notably, the two basidiomycete mycelia with the highest protein content were comparable to *R. microsporus* myc (**Fig. 3**). The crude protein content of *R. microsporus* (35.9 g/100 g DW) aligns with reported its value of 49.7 g/100 g DW, when accounting for differences in nitrogen-to-protein conversion factors (6.25 vs. 4.4 used in this study). *R. microsporus* myc represents the average crude protein content in mycelia of conventional fungi well (*Aspergillus oryzae* 45.7 g/100 g DW (Jin et al., 2002)) and total amino acid levels in *R. microsporus* myc are similar to *A. oryzae*, *Neurospora intermedia,* and *Fusarium venenatum*, and higher than *Rhizopus delemar (*Wang et al., 2023). Given that the crude protein content of *S. rugosa-annulata* and *S. commune* myc matched that of *R. microsporus*, these findings suggest that the protein content of basidiomycetous mycelia can be similar to CFF.

Protein quality is another important factor determining the suitability of mycelia to meet protein demands (Adhikari et al., 2022). In this study we analysed the protein quality of mycelium and fruiting bodies through the protein digestibility corrected amino acid score (PDCAAS), based on *in vitro* protein digestibility (IVPD) and amino acid composition. All samples had a relatively high IVPD, but basidiomycetous myc was significantly better digestible than their corresponding fruiting bodies, despite high variation between species (Supplementary fig. 6). This is in line with (Wallis et al., 2012), who observed a large variation in the protein digestibility between mushroom species, with most being suboptimal. Additionally, all basidiomycetous mycelia were significantly more digestible than *R. microsporus*, which has comparable digestibility, expressed as degree of hydrolysis, as other mycelia conventionally used in food, such as F. venenatum (R. Wang et al., 2023).

The nutritional quality of the amino acid composition, expressed by the amino acid score (AAS), was higher in basidiomycetous myc than in fruiting bodies and *R. microsporus* myc (Supplementary table 3). Six basidiomycetous mycelia contained complete amino acid profiles, except *V. volvacea* and *S. rugosa-annulata* mycelia, which were deficient in SAA and tryptophan (Trp), respectively. Low AAS for SAA were also found mycelial biomass (Wang et al., 2023) (Trp not measured). On the contrary, *R. microsporus* myc was deficient in both leucine and tryptophan. The combined nutritional effect of amino acid composition and IVPD, defined in the PDCAAS, was lowest in fruiting bodies (0.60 – 0.95), which was slightly higher than in most cereals (0.46 - 0.59), but lower than legumes (0.78 - 0.99) (Ertl et al., 2016; Zwinkels et al., 2023). Similarly, *R. microsporus* myc had a suboptimal PDCAAS (0.76), mainly due to the low IVPD. Nevertheless, the PDCAAS of all basidiomycetous mycelia was high (0.93 – 1.20), with four species surpassing a PDCAAS of 1.0, which is an indication of a complete protein. Thereby, the PDCAAS of basidiomycetous mycelia surpassed those of texturized soy-protein burgers (0.71 – 0.91) and were in the range of pork (1.17), beef (1.14), and chicken (0.92) (Ertl et al., 2016; Fanelli et al., 2022; Negrão et al., 2005). Moreover, lysine is the first limiting amino acid in most plant-based diets, making its PDCAAS crucial for assessing protein quality. Since soy is the primary plant-based lysine source but has limited scalability without further deforestation (Leinonen et al., 2019), diversifying non-animal lysine-rich proteins could significantly enhance global food security. Lysine was the limiting amino acid in two fruiting bodies, while in mycelia Trp, SAA, and Leu produced the lowest scores. Particularly in *S. commune* myc the PDCAAS of lysine was high (2.16), compared to *R. microsporus* (1.01) and fruiting bodies (0.60-1.01) (Supplementary fig. 8), and pork (1.62) (Fanelli et al., 2022).

The protein quality and protein content of mycelia and fruiting bodies were incorporated in the utilizable protein content, which is a measure to assess the amount of protein that can be absorbed by the body and utilized for protein synthesis and maintenance (Adhikari et al., 2023). All factors attributing to the utilizable protein content, namely a lower crude protein content, a lower IVPD and a less optimal amino acid composition, make fruiting bodies a suboptimal protein source. However, there is still a high interspecies variation, indicated by the almost double utilizable protein content in *L. edodes*, compared to *F. velutipes* and *C. comatus*. Similarly, the utilizable protein content of basidiomycetous mycelia varied greatly. The highest utilizable protein contents significantly exceeded that of *R. microsporus*. These results on protein content and quality indicate that there is a lot of variation between basidiomycetes, both between species and between vegetative and reproductive parts of the same species. However, the phylum of Basidiomycota contains several species of which the mycelium is a better source of protein than the commonly consumed filamentous fungus *R. microsporus*, which guides as a reliable benchmark for CFF in terms of protein content and quality. According to the FAO (2013), mycelium of *L. edodes*, *F. velutipes*, *S. commune*, and *P. cinnabarinus* can be considered “excellent” sources of protein, since they meet the criteria of a PDCAAS ≥ 1.0 and can provide ≥ 10.0 g crude protein per reference amount customarily consumed (estimated at 55 g per serving). In contrast with *R. microsporus* myc, which does not meet the protein quality criteria, and is therefore considered a “good” source of protein.

The selection of basidiomycetes studied only comprises a small portion of the enormous phylogenetic diversity in the phylum of Basidiomycota, which indicates much untapped potential. The results obtained suggest that mycelium of mushroom-producing fungi is a novel protein source with high potential, through combining a high umami taste with the high utilizable protein content found in mycelium of CFF. The high umami and aromatic content could make the mycelium palatable and gives it a recognizable taste profile. These results are in line with work by van Dam et al. (2024), who showed that mycelium from the basidiomycete *Pleurotus ostreatus* had a high consumer liking in a sensory panel. Safety of novel fungi is often a concern, due to potential secondary metabolite production, but according to Berger et al. (2022) there is little reason to assume that mycelia of edible basidiomycetes produce toxic compounds. One analogues source of proof for the safety of these mycelia is provided by van Dam et al. (2024), who observed that *P. ostreatus* myc was safe to consume, with lower mycotoxins in the mycelium than in the fruiting body. Nonetheless, only mycelium from shiitake (*L. edodes*) has been authorized for use under the EU novel food legislation under certain conditions (Turck et al., 2022). Therefore, new species would have to prove their safety by applying through this legislation.

Cultivation of these mycelia could take place in two main ways: (i) through submerged fermentation on food-grade substrates, pure mycelial biomass can be produced, similar to mycoprotein from *Fusarium venenatum*; (ii) in solid-state fermentation (SSF) on food-grade plant substrates, a mix of plant- and fungal material could be produced, similar to tempeh,. Recent work indicates that in both producing forms basidiomycetes can produce substantial amounts of fungal biomass, with similar efficiency to *R. microsporus* (Zwinkels et al., 2025, submitted (growth)). However, a longer fermentation time was required, reaching around eight days (in SSF), which is still considerably lower than in mushroom production, which generally takes weeks to months. The required usage of food-grade substrates limits the use of wood and hay-like substrates, typically used in mushroom production, but still allows to use food production side-streams, such as brewer’s spent grain, which has been fermented successfully already (Stoffel et al., 2019).

## 4 Materials and methods

### 4.1 Fungal species and mushrooms

*Lentinula edodes* CBS 134.85 (Shiitake), *Flammulina velutipes* CBS 347.74 (wild enoki), *Stropharia rugoso-annulata* CBS 288.85 (Garden giant), *Coprinus comatus* CBS 150.39 (Shaggy mane), *Schizophyllum commune* CBS 462.62 (Splitt gill fungus), *Volvariella volvacea* CBS 312.84 (Straw mushroom) and *Pycnoporus cinnabarinus* CBS 311.33 (Cinnabar polypore) and *Rhizopus microsporus* var. *oligosporus* CBS 338.62 were obtained from the public culture collection of Westerdijk Fungal Biodiversity Institute (Utrecht, The Netherlands). *Pleurotus pulmonarius* (M2345) (incorrectly portrayed by producer as *Pleurotus sajor-caju*) was obtained as mother culture on a petri dish from Mycelia BVBA (Deinze, Belgium). Basidiomycetes were stored on malt extract agar (MEA, Oxoid, Hampshire, United Kingdom) plates sealed with parafilm at 4°C, except for *V. volvacea* and *C. comatus*, which were stored at room temperature. *R. microsporus* was stored as a spore suspension in 20% (v/v) glycerol stock solution at −80°C.

for five species of basidiomycetes fruiting bodies were obtained. Shiitake mushrooms (dried whole, *L. edodes*) and enokitake (fresh, *F. velutipes*) were purchased from Amazing Oriental (Hoofddorp, The Netherlands). Wine cap mushrooms (fresh, *S. rugoso-annulata*) were purchased from Dennenhoeve Biologisch Dynamisch (Hooghalen, The Netherlands). Shaggy mane mushrooms (dried powder, *C. comatus*) were purchased from Naturaplaza.nl (Oldenzaal, The Netherlands). Paddy straw mushrooms (dried whole, *V. volvacea*) were obtained from SpiceJungle (Rockford, Michigan). All mushrooms were stored at −20°C until analysis.

### 4.2 Production process

#### 4.2.1 Basidiomycete and *Rhizopus* mycelia inoculum preparation

Basidiomycete inoculums were prepared according to the method described by Zwinkels et al. (2025, submitted), while *Rhizopus* inoculum was produced according to the method described by Wolkers – Rooijackers et al. (2018).

#### 4.2.2 Liquid culture fermentation

Mycelia were cultivated in 2 L conical shake flasks in 800 mL malt extract broth (MEB, Oxoid) containing tetracycline (25 mg/L, manufacturer, country). MEB was acidified using 1 M HCl to the optimal pH per fungal species derived from Zwinkels et al. (2025, submitted). Cultivation took place in the dark at the optimal temperature per species at 120 RPM until biomass growth plateaued visibly.

#### 4.2.3 Sample processing

Mycelia were filtered through wetted coffee filters and rinsed several times with demineralized water until the filtrate was clear. The rinsed mycelia were stored in 200 mL urine containers (VWR international, ML15) at −20°C. Mycelia and fruiting bodies were grounded into a fine powder by grinding for 30 s using a spice grinder (Krups F203, Solingen, Germany) after addition and evaporation of liquid nitrogen. Homogenised samples were lyophilized and again frozen at −20°C until further analysis.

### 4.3 Quantification of umami-active and volatile metabolites

#### 4.3.1 Free amino acids

Free amino acid concentrations in the mycelium and fruiting bodies were determined by following the method described by Scott et al. (2021).

#### 4.3.2 5’-nucleotides

5’-nucleotides (5-nuc) determination method was adapted from Seifar et al. (2009). Homogenized samples (100 mg) were dispersed in 5 mL milliQ water in a 15 mL centrifuge tube. Tubes were mixed by vortexing and heated for 2 min at 100°C in a shaking water bath. After heating, the tubes were vortexed for 30 s and cooled to room temperature. Samples were centrifuged (5 min, 10.000 ×*g*) and the supernatant was filtered using a Costar Spin-X centrifuge filter (0.22 µm, cellulose acetate). Subsequently, 200 µL filtered supernatant was stored in plastic autosample vials at −20°C until analysis. Separation, detection and quantification of 5’-nucleotides was performed using ultraperformance liquid chromatography (UPLC) with an UltiMate 3000 UPLC system (Dionex, Idstein, Germany) equipped with an AccQ-Tag Ultra BEH C_18_ column (150 mm x 2.1 mm, 1.7 µm) (Waters, Milford, MA, USA), and a BEH C_18_ guard column (5 mm x 2.1 mm, 1.7 µm) (Waters, Milford, MA, USA). The column temperature was set at 25°C and the mobile phase flow rate was maintained at 0.2 mL/min. Eluent A was composed of 10 mM dibutyl ammonium acetate (DBAA, pH 6.7) (TCI chemicals, Tokyo, Japan) and eluent B was composed of 10 mM DBAA (pH 6.7) containing 84% acetonitrile (Supelco, Bellefonte, PA, USA). The separation gradient was 0 – 2 min 99.9% A, 2 – 12 min steadily decreasing to 70% A, 12 – 15 min 70% A, 15 – 17 min 0% A, 17 – 27 min 99.9% A. One microliter of sample was injected for analysis and compounds were detected by UV measurements at 250 nm. Concentrations were calculated based on a calibration curve of 5’-adenosine monophosphate (Thermo Fisher), 5’-guanosine monophosphate (Thermo Fisher), 5’-inosine monophosphate (Thermo Fisher), and 5’-xanthosine monophosphate (Bio-Connect, Huissen, The Netherlands) (0.78 – 25 mg/L).

#### 4.3.3 Volatile organic compounds

Volatile organic compounds (VOCs) were determined by headspace solid phase microextraction gas chromatography mass spectrometry (HS SPME GC-MS) according to the method described by (Scott et al., 2021). In brief, a 0.1 gram homogenized and lyophilized mycelium and fruiting body sample was weighed in a 5 mL gas chromatography vial and capped. Vials were stored frozen (−20°C) until analysis. VOCs were detected in these ‘raw’ samples by HS SPME GC-MS. After the measurement, vials were heated by autoclaving them for 15 min at 121 °C. After this heating step, VOCs were again detected through HS SPME GC-MS under the same procedure (‘heated’ samples).

VOCs of mycelia and fruiting bodies were compared to meat. Ball tip steak (beef) and chicken breast filet were purchased fresh from Jumbo (Wageningen, The Netherlands). Samples were cut into 2 cm cubes and homogenized similarly to fungal materials. Subsequently, the dry matter content of grounded and homogenized meat samples was measured using a Moisture analyser (MA100, Sartorius). Using the dry matter content, two wet meat samples, equivalent to 0.1 g dry weight, were weighed in a GC vial. One of these samples was covered with aluminium foil and heated a convection oven at 150°C for 5 min. Both the ‘raw’ and ‘heated’ samples were stored frozen until analysis by HS SPME GC-MS. Only compounds with a MS Quantitation peak area over 20,000 counts*min were identified as detected.

### 4.4 Protein content and quality

#### 4.4.1 Dry matter content

Dry matter content was calculated from lyophilized samples (0.1 g) dried to a constant weight in a moisture analyser MA100 (Sartorius, Germany) at 105 °C. following the procedures described by the Association of Official Agricultural Chemists (AOAC, 2005).

#### 4.4.2 Crude protein content

Crude protein content was determined by DUMAS method with FlashSmart™ N/PROTEIN Element analyser (Thermo Scientific) in accordance with AOAC, 2006.03 (2006). Protein contents were calculated using food-specific nitrogen-to-protein conversion factors proposed by Mariotti et al. (2008). The used conversion factor for mycelium and fruiting bodies was 4.4.

#### 4.4.3 *In vitro* protein digestibility

*In vitro* protein digestibility (IVPD) was determined using the Megazyme testkit K-PDCAAS by following the manufacturer’s procedure with some adaptions (Megazyme, 2019). In brief, approximately 100 mg homogenized test, blank, control and standard sample was weighed into a 50 mL centrifuge tube, 19 mL HCl (0.06 M) was added, and tubes were incubated in a hot-air shaking incubator (37°C, 30 min, 300 RPM). Subsequently, proteins were digested with pepsin (60 min, 37°C, 300 RPM) after which samples were neutralized to pH 7.4 by addition of 2 mL 1.0 M Tris buffer (pH 7.4). Then, 200 µL trypsin/chymotrypsin mixture was added for the second digestion and samples were incubated for 4 h at 37°C at 300 RPM. This was followed by heating the samples in a boiling water bath for 10 min. Proteins were precipitated overnight at 4°C by addition of 1 mL TCA solution (40%). Tubes were centrifuged and subsequently the supernatants were diluted 10-fold and 20-fold in acetate buffer (50 mM, pH 5.5). 0.05 mL ninhydrin (2%) was added to 0.1 mL samples and incubated at 70°C, followed by through colorimetric determination of the amine concentration at 570 nm. L-glycine was used as calibration curve (0-1 mM). IVPD was calculated from the primary amine concentration of test samples fitted against standard samples with known *in vivo* protein digestibility values.

#### 4.4.4 Amino acid profile

The contents of amino acids were determined according to ISO13904 (ISO, 2005b) and ISO13903 (ISO, 2005a) procedures. In brief, to determine aromatic amino acids, phenylalanine and tyrosine, samples underwent acid hydrolysis. However, to determine tryptophan, samples underwent alkaline hydrolysis. All other amino acids were determined by oxidation before hydrolysis of the protein. The amino acids, except tryptophan were separated by ion exchange chromatography and determined by reaction with ninhydrin, using photometric detection at 570 nm and 440 nm for proline. Tryptophan was determined by reversed-phase C18 HPLC with fluorescence detection. These chemical analyses were executed in duplicate, and when the coefficient of variation was >5%, analyses were repeated.

#### 4.4.5 Protein quality

Protein quality was determined based on the protein digestibility corrected amino acid score (PDCAAS), in accordance with the FAO/WHO (1991). The amino acid score (AAS) was calculated using Equation 1. The reference protein was based on the dietary reference intake of a pre-school child (0.5– 3 y), as prescribed by the FAO (2013).

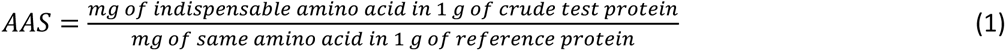

The PDCAAS was calculated by multiplying the IVPD by the AAS of the limiting amino acid (Equation 2).

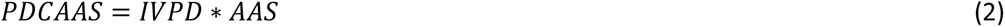

Adaptions to the calculations of PDCAAS were made to be more in line the more recently introduced digestible indispensable amino acid score (DIAAS). Namely, the PDCAAS exceeding 1.0 were not truncated to 1.0.

### 4.5 Data analysis

Data generated by UPLC and HS SPME GC-MS were analysed in Chromeleon 7.3.1 (Thermo Scientific, MA, USA). GC-MS peak integration was performed using the ICIS algorithm and the NIST main library was used for identification by matching mass spectral profiles with the profiles in NIST. One quantifying peak (generally the highest *m/z* peak per compound) was used per compound for quantification, while two confirming peaks were used for confirmation. Data were stored using Excel software v.16.0 (Microsoft Corporation, Redmond, WA, USA), data was analysed, and figures were produced using Rstudio software v. 4.0.2 (RStudio®, Boston, MA, USA). All values presented are means of biological triplicates ± standard deviation, unless stated otherwise. Significant differences that are indicated with letters were determined using One-Way ANOVA with Tukey post hoc test. Significance level (*p*-value) was set at 0.05.

## Supporting information

Supplementary file Zwinkels et al. 2025

## 7 Author contribution

The corresponding authors attest that all listed authors meet authorship criteria and that no others meeting the criteria have been omitted.

