## Supplementary file Zwinkels et al. 2025 for "Mycelium of mushroom-producing fungi as high-quality protein source"

### 1 Supplementary material

#### 1.1 Supplementary figures

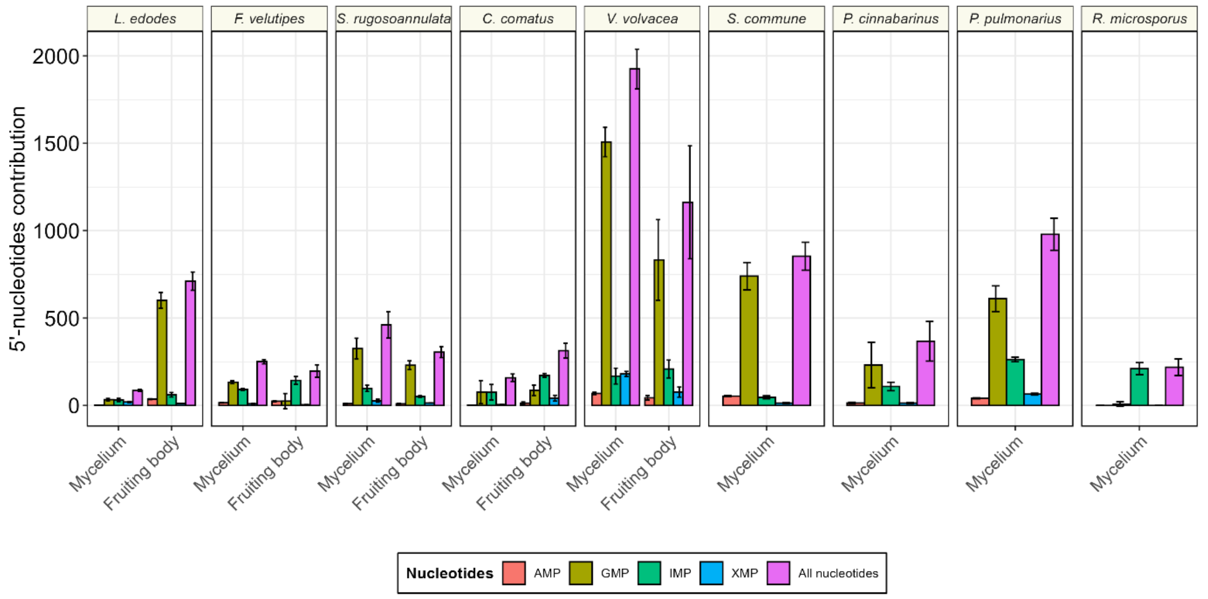

**Supplementary figure 1 | 5'-GMP is the main contributor to umami content.** Contribution of 5'-nucleotides to the equivalent umami concentration of mycelia and fruiting bodies. Colours represent the 5'-nucleotides, 5'-adenosine monophosphate (AMP, red), 5'-guanosine monophosphate (GMP, olive green), 5'-inosine monophosphate (IMP, green), 5'-xanthanosine monophosphate (XMP, blue), and the sum of all nucleotides contributions (purple).

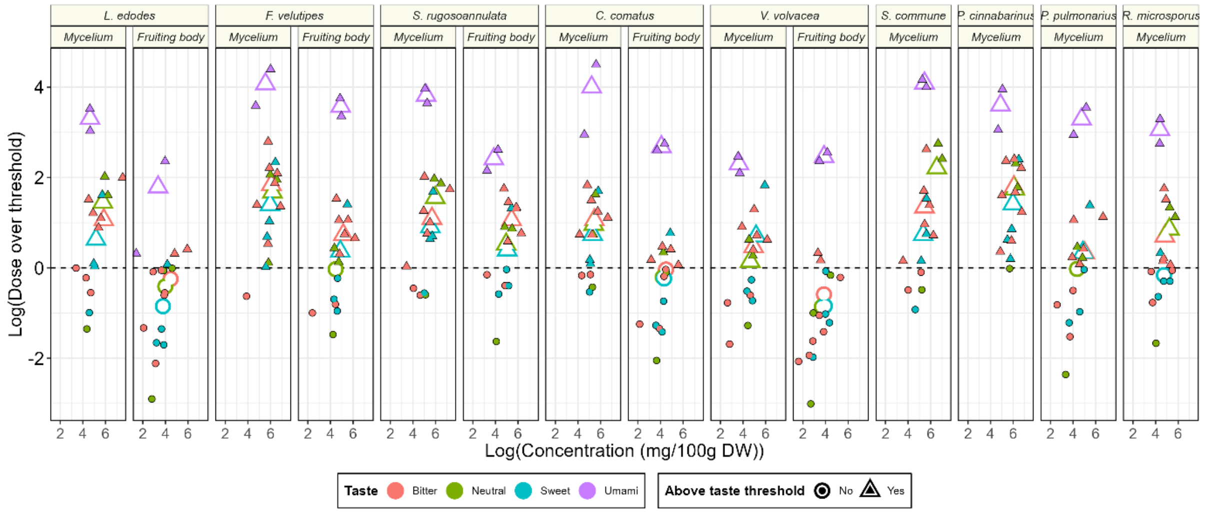

**Supplementary figure 2 | Dose over threshold is consistently highest for umami free amino acids.** Relationship between log(Dose over threshold) and log(concentration (g/100 g DW) of free amino acids with bitter (red), neutral (green), sweet (blue), or umami (purple) taste in mycelia and fruiting bodies. Closed points represent individual amino acids, while open points represent the average of all amino acids in a certain taste category. Round points are below the taste threshold, while triangular points are above the taste threshold.

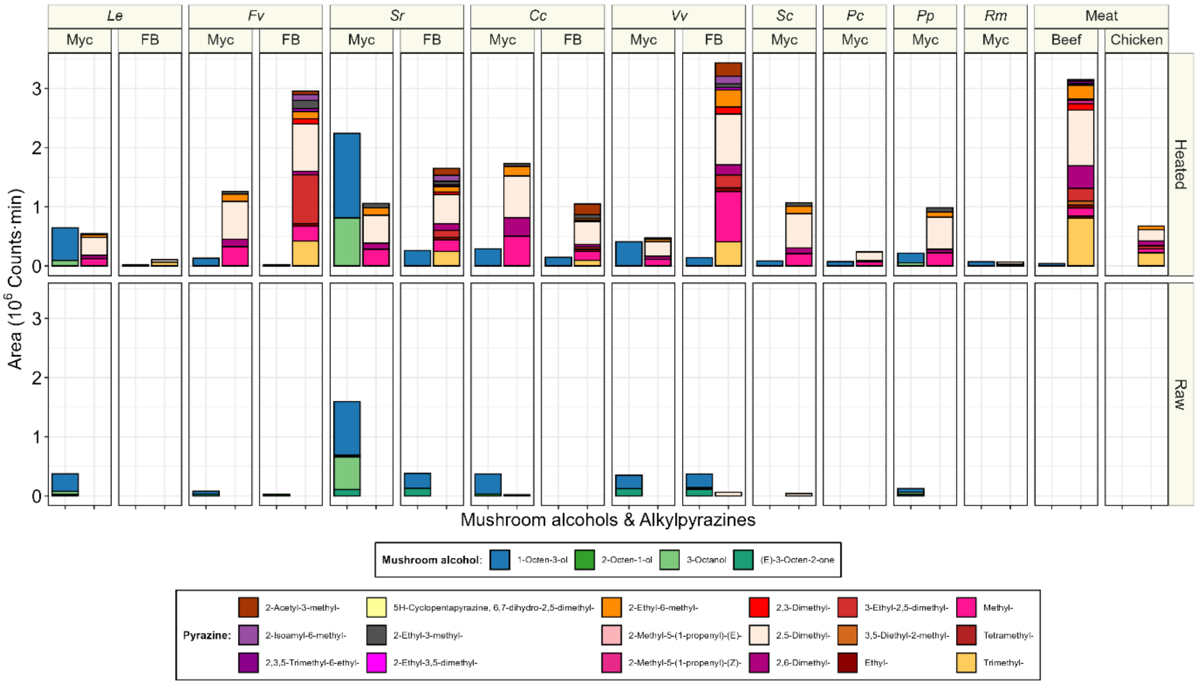

**Supplementary figure 3 | Alkylpyrazines are formed during heating both in mycelia and fruiting bodies.** Mushroom alcohol (left) and alkylpyrazine (right) area ($10^6$ counts*min) in raw and heated mycelium (Myc) and fruiting bodies (FB), chicken and beef. Species are *L. edodes* (*Le*), *F. velutipes* (*Fv*), *S. rugosa-annulata* (*Sr*), *C. comatus* (*Cc*), *V. volvacea* (*Vv*), *S. commune* (*Sc*), *P. cinnabarinus* (*Pc*), *P. pulmonarius* (*Pp*), and *R. microsporus* (*Rm*). Colours represent compounds and are explained in the figure legend.

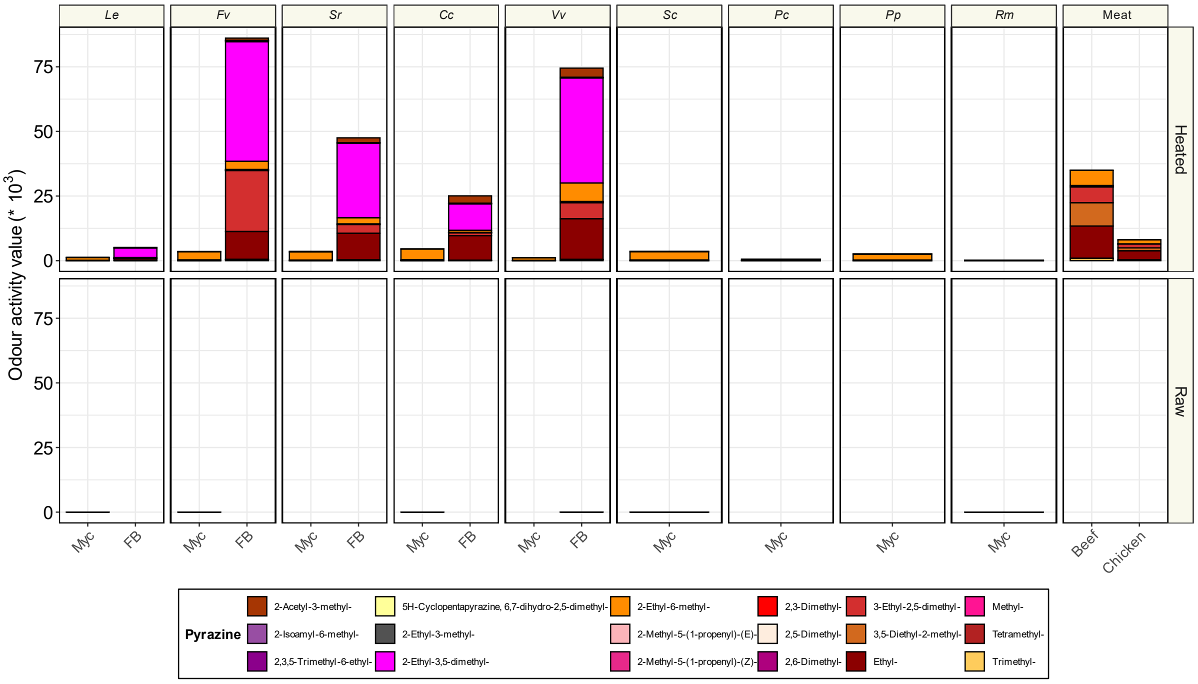

**Supplementary figure 4 | Odour activity (OAV) value is largest in fruiting bodies.** OAV of alkylpyrazines (*$10^3$) in raw and heated mycelium (Myc) and fruiting bodies (FB), chicken and beef. OAV is calculated as: Area (counts*min)/Odour threshold (µg/L). Odour thresholds are based on literature and provided in Supplementary table 1. Species are *L. edodes* (*Le*), *F. velutipes* (*Fv*), *S. rugosa-annulata* (*Sr*), *C. comatus* (*Cc*), *V. volvacea* (*Vv*), *S. commune* (*Sc*), *P. cinnabarinus* (*Pc*), *P. pulmonarius* (*Pp*), and *R. microsporus* (*Rm*). Colours represent compounds and are explained in the figure legend.

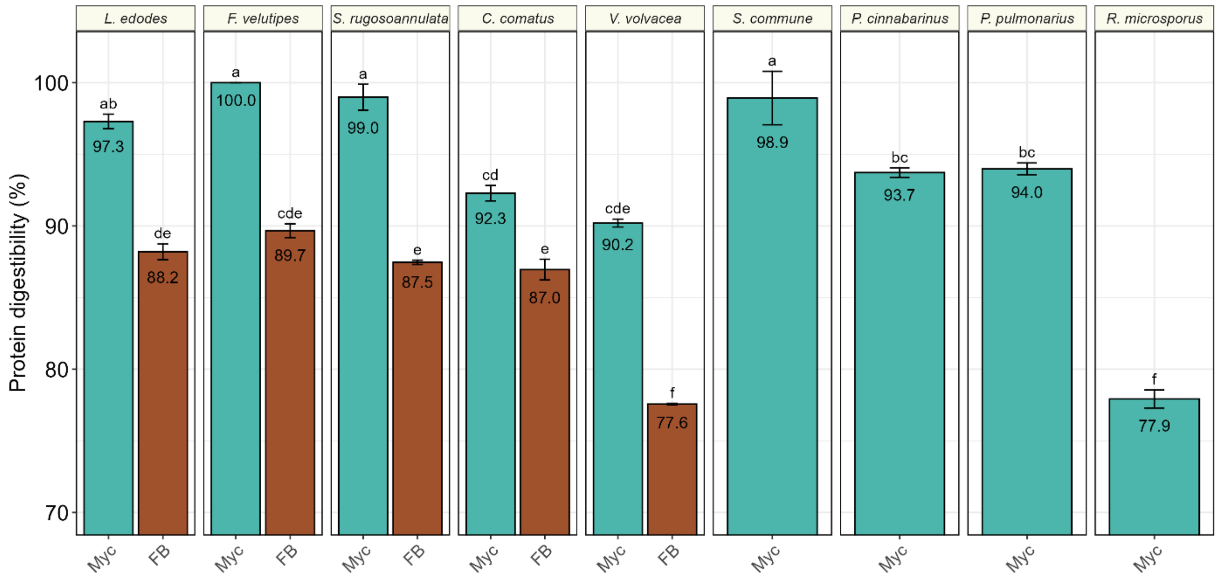

**Supplementary figure 5 | *In vitro* protein digestibility (%) of mycelia (cyan) and fruiting bodies (brown).** Bars are means of biological triplicates ± standard deviation. Numbers indicate the mean crude protein content. Different letters indicate a significant difference (TukeyHSD, p<0.05).

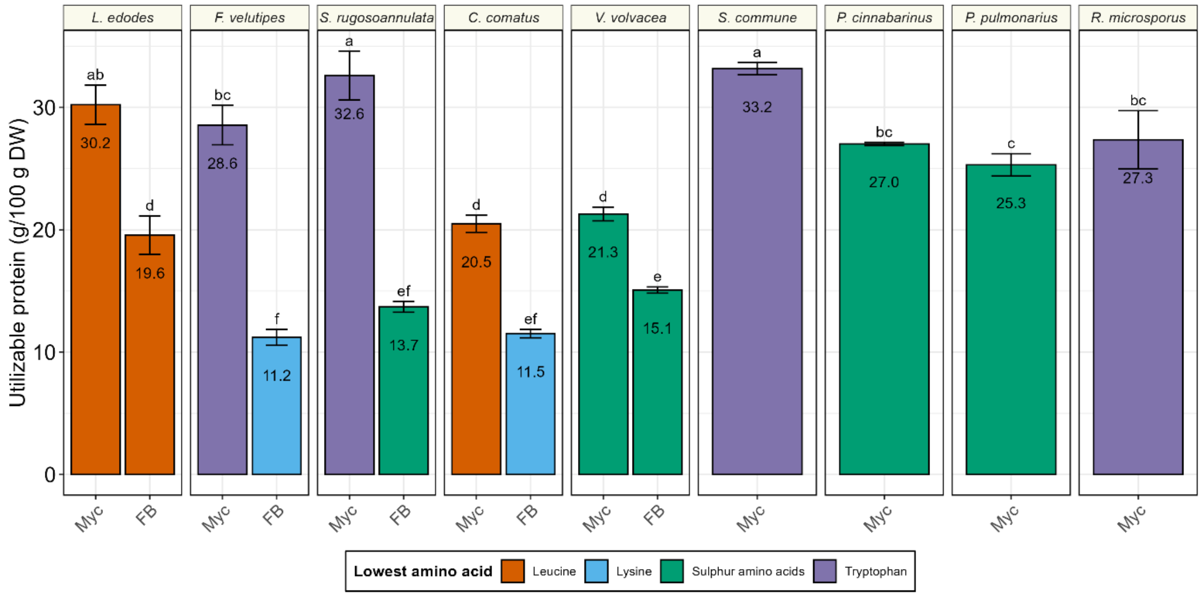

**Supplementary figure 6 | Utilizable protein content (g/100 g DW) of mycelia and fruiting bodies.** Colours indicate the lowest amino acids leucine (brown), lysine (light blue), sulphur amino acids (SAA, green), and tryptophan (purple). Bars are means of biological triplicates ± standard deviation. Numbers indicate the mean crude protein content. Different letters indicate a significant difference (TukeyHSD p<0.05).

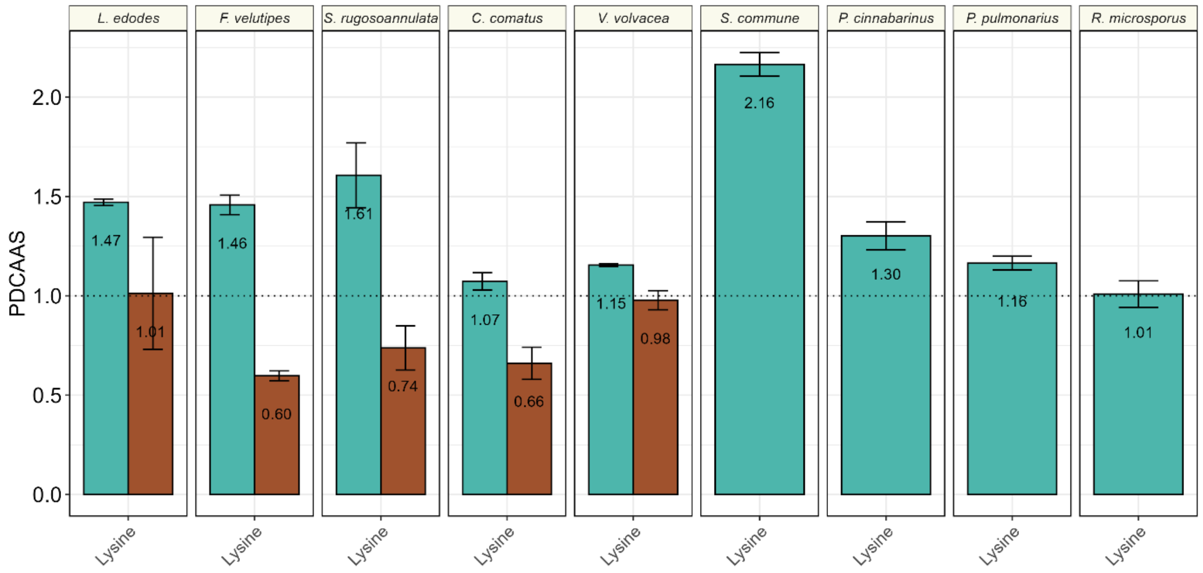

**Supplementary figure 7 | Protein digestibility corrected amino acid score (PDCAAS) of mycelium and fruiting body of basidiomycetes and *Rhizopus*.** Bars are means of biological triplicates ± standard deviation. Colours indicate the lowest amino acids leucine (red), lysine (green), sulphur amino acids (SAA, blue), and tryptophan (purple). Lowest amino acid is the limiting amino acid when PDCAAS < 1.0 (dotted line).

#### 1.2 Supplementary tables

**Supplementary table 1.** Odor description and odour threshold in water (µg/L) of alkylpyrazines present in mycelia, fruiting bodies or meat (Cherniienko et al., 2022; Shi et al., 2025; Sohail et al., 2022; Yan et al., 2021).

| Alkylpyrazines | Odour description | Odor threshold range in water (in µg/L) | Odor threshold used in OAV |
|---|---|---|---|
| 2,3,5-Trimethyl-6-ethylpyrazine | chocolate, cocoa, coffee, sweet, hazelnut, roasted | 0,002-0,024 | 0.013 |
| 2,3-Dimethylpyrazine | nutty, cocoa, peanut butter, coffee, caramel, roasted potato, musty | 400-2500 | 1450 |
| 2,5-Dimethylpyrazine | cocoa, roasted, nutty, beef, woody, grassy, medicinal, earthy | 1700-2600 | 2150 |
| 2,6-Dimethylpyrazine | ether, cocoa, nutty, roasted, beefy, coffee, buttermilk | 400-9000 | 4700 |
| 2-Acetyl-3-methylpyrazine | nutty, popcorn, roasted corn, dirt, burnt, sweet | 62 | 62 |
| 2-Ethyl-3,5-dimethylpyrazine | burnt, coffee, nutty, roasted, woody, potato-chip-like, | 0,04-2 | 1.02 |
| 2-Ethyl-3-methylpyrazine | nutty, musty, corn, raw, earthy, bready | 130-500 | 315 |
| 2-Ethyl-6-methylpyrazine | roasted, potato | 40 | 40 |
| 2-Methyl-5-(1-propenyl)-(E)-pyrazine | Roasted, nutty | - | - |
| 2-Methyl-5-(1-propenyl)-(Z)-pyrazine | Roasted, nutty | - | - |
| 3,5-Diethyl-2-methylpyrazine | nutty, meaty, vegetable | 7.5 | 7.5 |
| 3,5-Dimethyl-2-isobutylpyrazine | roasted, sweet, green | - | - |
| 3-Ethyl-2,5-dimethylpyrazine | nutty, roasted sunflower seeds | 35 | 35 |
| 6,7-Dihydro-2,5-dimethyl-5H-cyclopentapyrazine | - | - | - |
| Methylpyrazine | pungent, sweet, corn-like, nutty, chocolate, hazelnut, green | 27.000-100.000 | 63500 |
| Tetramethylpyrazine | nutty, musty, chocolate, coffee, cocoa, burnt, musty, vanilla | 1000-38.000 | 19500 |
| Trimethylpyrazine | nutty, earthy, powdery, cocoa, potato, roasted | 90-1800 | 945 |

**Supplementary table 2 | Amino acid content (mg/g DW) and indispensable amino acid index (IAAI) of mycelia and fruiting bodies.** Values are means of biological triplicates ± standard deviation. Myc = mycelium, FB = Fruiting body

| | ***L. edodes*** | | | | ***F. velutipes*** | | | | ***S. rugoso-annulata*** | | | | ***C. comatus*** | | | | ***V. volvacea*** | | | | ***S. commune*** | | ***P. cinnabarinus*** | | ***P. pulmonarius*** | | ***R. microsporus*** | |
|---|---|---|---|---|---|---|---|---|---|---|---|---|---|---|---|---|---|---|---|---|---|---|---|---|---|---|---|---|
| Amino acid | **Myc** | | **FB** | | **Myc** | | **FB** | | **Myc** | | **FB** | | **Myc** | | **FB** | | **Myc** | | **FB** | | **Myc** | | **Myc** | | **Myc** | | **Myc** | |
| (mg/g DW) | **mean** | **sd** | **mean** | **sd** | **mean** | **sd** | **mean** | **sd** | **mean** | **sd** | **mean** | **sd** | **mean** | **sd** | **mean** | **sd** | **mean** | **sd** | **mean** | **sd** | **mean** | **sd** | **mean** | **sd** | **mean** | **sd** | **mean** | **sd** |
| Alanine | 20,49 | 2,62 | 12,3 | 1,65 | 25,22 | 0,52 | 10,63 | 0,79 | 26,03 | 1,41 | 10,74 | 0,4 | 15,77 | 0,33 | 9,68 | 0,39 | 22 | 0,68 | 16,61 | 0,21 | 25,18 | 0,43 | 24,14 | 1,96 | 18,37 | 0,66 | 19,94 | 1,47 |
| Arginine | 31,79 | 1,91 | 12,7 | 2,63 | 20,07 | 1,23 | 6,42 | 0,38 | 29,34 | 0,56 | 6,06 | 0,28 | 12,47 | 0,75 | 9,81 | 0,18 | 18,63 | 0,57 | 13,4 | 0,32 | 25,55 | 1,25 | 20,96 | 0,97 | 22,93 | 1,11 | 16,02 | 1,08 |
| Aspartic acid | 30,52 | 2,75 | 22,01 | 3,32 | 27,81 | 0,82 | 11,1 | 0,52 | 38,49 | 2,4 | 13,9 | 0,55 | 21,99 | 0,48 | 14,21 | 0,53 | 27,18 | 0,73 | 25,46 | 0,55 | 32,2 | 0,62 | 29,73 | 1,25 | 24,77 | 0,81 | 33,37 | 3,35 |
| Cysteine | N.D. | N.D. | N.D. | N.D. | 2,96 | 0,11 | N.D. | N.D. | 5,54 | 0,35 | 2,02 | 0,07 | 3,15 | 0,06 | N.D. | N.D. | 2,4 | 0,15 | 2,3 | 0,04 | 4,02 | 0,09 | 3,95 | 0,11 | 3,08 | 0,09 | 5,45 | 0,81 |
| Glutamic acid | 41,79 | 1,71 | 42,21 | 17,57 | 48,87 | 1,32 | 14,17 | 0,49 | 58,19 | 3,18 | 25,13 | 1,01 | 33,78 | 0,83 | 17,05 | 0,6 | 33,47 | 2,76 | 43,54 | 1,11 | 44,47 | 0,15 | 42,18 | 0,83 | 42,01 | 3,25 | 39,1 | 3,29 |
| Glycine | 16,05 | 2,55 | 10,68 | 1,96 | 17,94 | 0,57 | 7,18 | 0,14 | 21,67 | 0,61 | 7,87 | 0,36 | 12,05 | 0,21 | 6,87 | 0,71 | 13,98 | 0,34 | 11,79 | 0,17 | 19,06 | 1,38 | 16,42 | 0,75 | 13,17 | 0,39 | 15,35 | 1,33 |
| Histidine | 8,26 | 0,7 | 5,33 | 0,9 | 9,76 | 0,41 | 4,02 | 0,19 | 10,65 | 0,74 | 4,18 | 0,19 | 5,52 | 0,16 | 3,4 | 0,11 | 6,86 | 0,12 | 5,53 | 0,05 | 10,82 | 0,97 | 8,6 | 0,4 | 7,03 | 0,29 | 10,79 | 0,81 |
| Isoleucine | 15,37 | 1,54 | 10,01 | 1,49 | 14,69 | 0,53 | 6,06 | 0,32 | 17,35 | 1,1 | 6,81 | 0,21 | 9,74 | 0,35 | 6,37 | 0,19 | 14,07 | 0,07 | 11,93 | 0,25 | 16,13 | 0,47 | 13,69 | 0,76 | 11,56 | 0,49 | 15,95 | 1,12 |
| Leucine | 23,78 | 2,51 | 15,41 | 2,34 | 23,13 | 0,87 | 9,18 | 0,45 | 27,64 | 1,35 | 11,24 | 0,36 | 14,65 | 0,5 | 9,35 | 0,27 | 22,18 | 0,47 | 19,17 | 0,41 | 26,59 | 0,63 | 23,81 | 1,17 | 18,97 | 0,68 | 23,42 | 1,51 |
| Lysine | 26,03 | 1,19 | 14,14 | 3,44 | 23,7 | 0,76 | 7,12 | 0,4 | 32,43 | 3,93 | 9,86 | 0,43 | 14,54 | 0,28 | 7,55 | 0,29 | 18,97 | 0,44 | 16,61 | 0,3 | 41,36 | 0,56 | 21,39 | 1,2 | 18,22 | 0,72 | 26,36 | 1,39 |
| Methionine | N.D. | N.D. | N.D. | N.D. | 5,33 | 0,13 | N.D. | N.D. | 5,46 | 0,27 | 2,21 | 0,07 | 3,59 | 0,09 | N.D. | N.D. | 3,98 | 0,03 | 2,95 | 0,04 | 6,23 | 0,22 | 5,42 | 0,32 | 4,2 | 0,16 | 5,77 | 0,39 |
| Phenylalanine | 14,38 | 1,22 | 9,67 | 1,5 | 13,56 | 0,64 | 7,67 | 0,46 | 14,27 | 0,96 | 7,87 | 0,19 | 7,92 | 0,33 | 5,78 | 0,19 | 11,87 | 0,39 | 10,86 | 0,84 | 13,3 | 2,02 | 12,54 | 0,95 | 11,01 | 0,73 | 13,95 | 1,52 |
| Proline | 13,92 | 1,93 | 8,4 | 1,26 | 15,61 | 0,37 | 5,18 | 0,09 | 17,64 | 0,2 | 6,49 | 0,26 | 9,59 | 0,13 | 5,8 | 0,17 | 13,66 | 0,16 | 10,93 | 0,12 | 16,84 | 1,09 | 15,3 | 0,64 | 11,55 | 0,24 | 12,86 | 0,96 |
| Serine | 15,71 | 1,36 | 10,56 | 1,51 | 16,4 | 0,61 | 6,03 | 0,41 | 21,9 | 1,12 | 8,37 | 0,37 | 11,21 | 0,35 | 7,82 | 1,29 | 14,24 | 0,23 | 14,05 | 0,25 | 16,85 | 0,13 | 17,32 | 0,55 | 14,38 | 0,51 | 15,73 | 1,36 |
| Threonine | 15,37 | 1,13 | 10,52 | 1,86 | 16,42 | 0,5 | 7,06 | 0,11 | 19,42 | 1 | 8,47 | 0,25 | 11,46 | 0,36 | 7,34 | 0,54 | 15,46 | 0,58 | 13,44 | 0,17 | 16,41 | 0,45 | 16,42 | 0,5 | 13,57 | 0,71 | 16,46 | 1,43 |
| Tryptophan | N.D. | N.D. | N.D. | N.D. | 2,56 | 0,24 | N.D. | N.D. | 2,8 | 0,16 | 1,61 | 0,04 | 2,02 | 0,05 | N.D. | N.D. | 2,97 | 0,03 | 2,73 | 0,06 | 3,04 | 0,07 | 3,01 | 0,07 | 2,42 | 0,08 | 2,98 | 0,26 |
| Tyrosine | 7,37 | 0,76 | 2,62 | 0,68 | 6,69 | 0,9 | 3,81 | 0,19 | 7,21 | 0,7 | 3,2 | 0,36 | 4,78 | 0,19 | 2,57 | 0,1 | 6,85 | 0,41 | 4,28 | 0,42 | 6,85 | 1,75 | 6,86 | 0,48 | 5,82 | 0,4 | 8,36 | 1,17 |
| Valine | 17,34 | 2,02 | 11,63 | 1,83 | 17,74 | 0,6 | 7,35 | 0,42 | 20,87 | 1,11 | 8,51 | 0,37 | 13,45 | 0,47 | 7,89 | 0,25 | 16,44 | 0,26 | 14,08 | 0,28 | 20,22 | 0,36 | 16,99 | 0,84 | 13,92 | 0,46 | 17,56 | 1,27 |
| IAA[a] | 120,53 | 10,03 | 76,72 | 13,33 | 126,9 | 4,54 | 48,46 | 2,04 | 150,88 | 10,19 | 60,77 | 1,75 | 82,87 | 1,59 | 47,69 | 1,76 | 112,79 | 0,34 | 97,3 | 1,92 | 154,1 | 5,02 | 121,88 | 5,84 | 100,9 | 4,16 | 133,25 | 9,44 |
| DAA[b] | 179,08 | 14,6 | 123,15 | 29,61 | 181,55 | 5,75 | 68,56 | 3,06 | 226,02 | 8,93 | 83,77 | 3,41 | 124,79 | 2,51 | 76,05 | 4,1 | 152,4 | 3,51 | 142,36 | 2,86 | 191,03 | 0,49 | 176,84 | 6,59 | 156,06 | 7,36 | 166,19 | 13,6 |
| IAAI[c] | 0,40 | 0 | 0,39 | 0,02 | 0,41 | 0 | 0,41 | 0 | 0,40 | 0,01 | 0,42 | 0 | 0,40 | 0 | 0,39 | 0 | 0,43 | 0 | 0,41 | 0 | 0,45 | 0,01 | 0,41 | 0 | 0,39 | 0 | 0,45 | 0 |

[a] Indispensable amino acids
[b]Dispensable amino acids
[c]Indispensable amino acid index
N.D. = Not Determined.

**Supplementary table 3 | Amino acid content of indispensable amino acids (mg/g crude protein),** presented as means of biological triplicates ± standard deviation, compared to the dietary requirements of infants (0-6 months), children (6 months – 3 year), and adults (>3 year). Amino acid contents not meeting dietary requirements of children (6m-3y) are marked in red.

| | | Aromatic amino acids | | Histidine | | Isoleucine | | Leucine | | Lysine | | Sulphur amino acids | | Threonine | | Tryptophan | | Valine | |
|---|---|---|---|---|---|---|---|---|---|---|---|---|---|---|---|---|---|---|---|
| **Species** | **Material** | **mean** | **sd** | **mean** | **sd** | **mean** | **sd** | **mean** | **sd** | **mean** | **sd** | **mean** | **sd** | **mean** | **sd** | **mean** | **sd** | **mean** | **sd** |
| *L. edodes* | Mycelium | 72,0 | 4,3 | 27,3 | 1,9 | 50,8 | 4,2 | 78,6 | 6,8 | 86,2 | 1,0 | N.D. | | 50,8 | 2,5 | N.D. | | 57,4 | 5,7 |
| *L. edodes* | Fruiting body | 56,7 | 6,1 | 24,6 | 4,9 | 46,2 | 8,2 | 71,2 | 12,8 | 65,4 | 17,8 | N.D. | | 48,6 | 10,0 | N.D. | | 53,7 | 10,0 |
| *F. velutipes* | Mycelium | 70,9 | 1,8 | 34,2 | 0,5 | 51,5 | 1,3 | 81,1 | 1,9 | 83,1 | 2,8 | 29,1 | 1,2 | 57,6 | 1,7 | 9,0 | 0,3 | 62,2 | 1,5 |
| *F. velutipes* | Fruiting body | 61,3 | 3,3 | 21,5 | 2,5 | 32,3 | 1,4 | 49,0 | 2,3 | 38,0 | 1,5 | N.D. | | 37,7 | 3,0 | N.D. | | 39,2 | 1,6 |
| *S. rugoso-annulata* | Mycelium | 61,4 | 4,6 | 30,4 | 1,3 | 49,5 | 2,2 | 78,9 | 2,6 | 92,5 | 8,9 | 31,4 | 1,5 | 55,5 | 2,0 | 8,0 | 0,3 | 59,6 | 2,1 |
| *S. rugoso-annulata* | Fruiting body | 53,8 | 5,1 | 20,4 | 2,9 | 33,2 | 4,9 | 54,8 | 8,1 | 48,1 | 7,2 | 20,6 | 2,8 | 41,3 | 5,9 | 7,9 | 1,2 | 41,5 | 6,3 |
| *C. comatus* | Mycelium | 57,8 | 3,2 | 25,1 | 0,3 | 44,3 | 0,9 | 66,7 | 1,5 | 66,2 | 2,3 | 30,7 | 0,9 | 52,2 | 1,1 | 9,2 | 0,2 | 61,2 | 1,2 |
| *C. comatus* | Fruiting body | 47,9 | 5,6 | 19,5 | 2,8 | 36,5 | 4,4 | 53,6 | 6,6 | 43,2 | 4,9 | N.D. | | 41,9 | 3,3 | N.D. | | 45,2 | 5,5 |
| *V. volvacea* | Mycelium | 72,1 | 4,5 | 26,4 | 0,3 | 54,1 | 1,3 | 85,4 | 1,3 | 73,0 | 0,6 | 24,5 | 0,5 | 59,5 | 3,4 | 11,4 | 0,2 | 63,3 | 2,3 |
| *V. volvacea* | Fruiting body | 65,4 | 4,8 | 23,9 | 1,1 | 51,6 | 3,0 | 82,9 | 4,6 | 71,9 | 3,5 | 22,7 | 1,2 | 58,1 | 2,6 | 11,8 | 0,7 | 60,9 | 3,3 |
| *S. commune* | Mycelium | 60,7 | 10,8 | 32,6 | 2,4 | 48,6 | 1,1 | 80,2 | 1,3 | 124,7 | 1,2 | 30,9 | 0,6 | 49,5 | 0,8 | 9,2 | 0,1 | 61,0 | 1,0 |
| *P. cinnabarinus* | Mycelium | 71,8 | 4,9 | 31,8 | 1,4 | 50,7 | 2,6 | 88,1 | 3,9 | 79,2 | 4,1 | 34,7 | 1,4 | 60,8 | 1,6 | 11,2 | 0,3 | 62,9 | 2,8 |
| *P. pulmonarius* | Mycelium | 65,2 | 3,3 | 27,2 | 0,7 | 44,8 | 1,6 | 73,6 | 2,2 | 70,6 | 1,9 | 28,2 | 0,8 | 52,6 | 2,2 | 9,4 | 0,2 | 54,0 | 1,4 |
| *R. microsporus* | Mycelium | 62,2 | 3,4 | 30,2 | 1,1 | 44,6 | 1,5 | 65,5 | 2,5 | 73,8 | 4,4 | 31,3 | 0,5 | 45,9 | 0,9 | 8,3 | 0,2 | 49,0 | 1,5 |
| Dietary requirements[a] | Infants (0-6m) | 94 | | 21 | | 55 | | 96 | | 69 | | 33 | | 44 | | 17 | | 55 | |
| Dietary requirements[a] | Children (6m-3y) | 52 | | 20 | | 32 | | 66 | | 57 | | 27 | | 31 | | 8,5 | | 43 | |
| Dietary requirements[a] | Adults (≥3y) | 41 | | 16 | | 30 | | 61 | | 48 | | 23 | | 25 | | 6,6 | | 40 | |

[a]Based on dietary recommendations set by FAO (2013)
